# Computational Insights into the tACS Modulation in Healthy and Epileptic Brain Networks

**DOI:** 10.64898/2026.06.23.733961

**Authors:** Mariam Al Harrach, Maxime Yochum, Gabriel Gaugain, Julien Modolo, Fabrice Wendling

## Abstract

Transcranial Electric stimulation (tES) is a safe and noninvasive technique increasingly used in treating brain disorders. Despite many studies on tES, there is still a lack of understanding of its mechanisms at the microscale network level. This is crucial for optimized parameter selection in therapy approaches such as the treatment of pharmacoresistant epilepsy. In this study, we made use of a recently published neuroinspired microscale model of the neocortex, known as NeoCoMM, and integrated a “Lambda E”-based model of tES. This updated version was used to investigate the acute effects of tES (tDCS and tACS), on the neural activity of various neuron types in both healthy and epileptic brain states. Results showed that in the case of healthy alpha and gamma rhythms, tACS induced electric field entrainment at the peak power frequency of the network oscillations as measured by the Local field Potentials (LFPs). This resonance-like entrainment was independent from the individual firing rate of cell types. For epileptic activity, tACS did not provide consistent results. Cathodal tDCS resulted in a promising decrease in hyperexcitable activity throughout simulations. These results advance our understanding of the impact of tES on network dynamics at both the extracellular and intracellular activity levels.

**Author summary:** 

## Introduction

Cerebral activity is characterized by oscillations that result from interactions between interconnected excitatory (Glutamatergic) and inhibitory (GABAergic) neurons comprised in neural networks [1]. These oscillations are the basis of multiple functions, and irregularities in these rhythms are usually related to underlying psychiatric and neurological disorders [2, 3]. As such, neuromodulation techniques are being continuously investigated by both researchers and clinicians to uncover their ability to modulate brain oscillations in healthy and pathological circuits [4].

Transcranial Electric Stimulation (tES) is a neuromodulation technique that consists in applying electrical currents to the scalp. It is safe, non-invasive, well-tolerated, and compatible with pharmacotherapy, and it can even be used by patients at home [5]. Consequently, tES has drawn an increasing interest for its promising therapeutic potential in psychiatric and neurological disorders and for its utility in studying brain function in general [3, 6–9]. tES techniques include transcranial Alternating Current Stimulation (tACS) and transcranial Direct Current Stimulation (tDCS). While tACS targets neural oscillations to influence brain activity patterns, tDCS focuses on cortical excitability to either increase or decrease neuronal excitability [1].

In recent years, tES has gained increasing attention for its potential therapeutic applications in psychiatric and neurological disorders, and for its utility in studying brain function [6–8]. There has been a significant rise in clinical trials investigating tES as a promising clinical tool in depression [10, 11], dementia [12], dystonia [13], Parkinson’s Disease [14], and epilepsy [6, 15, 16]. Accordingly, understanding the underlying physiological and biophysical mechanisms of both tDCS and tACS has become of great interest to optimize their therapeutic potentials [17–21].

The effect of tDCS was initially believed to affect the membrane potential of cells, resulting in hyperpolarisation or depolarisation of the resting membrane potential for cathodal and anodal tDCS, respectively [16, 22]. However, studies have reported that tDCs may not only have a linear effect on the firing rate but also on neuronal transmission [23, 24]. In the case of tACS, early research emphasized straightforward effects, such as entraining neuronal oscillations to the applied alternating current. However, recent studies highlighted the complex, nonlinear interactions between the induced electrical fields and ongoing brain dynamics [25].

Despite these studies, our understanding of how tDCS and tACS affect neurons and neural networks is still incomplete. In particular, in the case of tACS, we have yet to fully investigate how varying stimulation strength and frequency impacts the different types of neurons comprised in brain circuits, as well as their interactions [9]. Indeed, the complexity is that there is evidence that tACS differentially impacts cellular types, independently of their connectivity [26]. In addition, connectivity adds further difficulty in disentangling tACS effects. Therefore, uncovering tACS underlying mechanisms can be an asset in generalizing the use and optimization of tES for neurological disorders.

Epilepsy is one of the predominant neurological diseases in the world [27, 28], affecting around 65 million people and being characterized by recurrent, spontaneous seizures [28]. About 36.3 % of epilepsy patients are drug-resistant [29], which calls for developing alternative therapeutic strategies [30, 31]. In this context, transcranial electrical stimulation (tES) is a potential approach for reducing seizure frequency and improving outcomes in these patients. tES has been shown to be safe and very promising [32, 33]; however, although some trials and clinical studies showed a reduction in seizure frequency with direct current stimulation (tDCS) [33, 34], the percentage of responders (about 50 percent of patients experiencing a 50 percent reduction in seizure frequency) could certainly be increased. One challenge is that uncovering tES effects on epileptic tissue in vivo involves complex mechanistic and feasibility issues, especially since the parameter space is high. This reinforces the need for computational modeling approaches to optimize the stimulation strategy by uncovering the underlying coupled neurobiological/biophysical mechanisms induced by tES.

Here, we propose a novel neuroinspired microscopic (celluar scale) model [35, 36] called NeoCoMM (NeoCortical Computational Microscale Model) that simulates the cortical column and that is used to study the acute effects of tES (tDCS and tACS) on healthy and epileptogenic neocortical neuronal networks. NeoCoMM is an open source model developed in our team [36] that allows for the simulation of realistic extracellular and intracellular activity of a network of neocortical neurons and interneurons. NeoCoMM can be configured to simulate healthy or pathological brain oscillations. In this work, we used this model and developed a new, expanded version of NeoCoMM that can simulate the impact of tACS and tDCS on microcircuits and provides both extracellular and intracellular activity of glutamatergic and GABAergic neurons comprised in the considered patch of neocortex tissue.

First, we considered normal brain rhythms that consisted of simulations in the alpha (8-12 Hz) and low gamma (30-50 Hz) frequency bands. We used these simulations to study the impact of tACS frequency and intensity on network and cell dynamics. We found that increased neuronal entrainment is obtained when stimulation is delivered at peak frequency of the spontaneous network activity. The significance of neuromodulation depends on network activity rather than cell-specific dynamics. We then investigated the impact of tACS frequency and intensity on interictal epileptic activity. This analysis did not reveal a potenial effect of tACS for decreasing brain tissue hyperexcitability and for decreasing epileptiform events.

Regarding simulated direct currents, results showed that cathodal tDCS at human therapy relevant intensities resulted in a decrease in the number of epileptiform events, bursting of excitatory cells, overall explained by a decrease of the ratio between excitatory and inhibitory processes. This result confirms recent findings regarding the promising efficacy of tDCS in the treatment of pharmacoresistant epilepsy [34].

Overall, our upgraded model brings a significant contribution by featuring the effects of in situ electric fields induced transcranially, providing a precious tool for the tES community. It can elucidate the underlying mechanistic impacts of tES protocols by simulating their impact on extracellular as well as cellular activity. Furthermore, while this upgraded version of NeoCoMM focuses on acute (immediate) effects, it provides a basis for future neuroplasticity models aimed at investigating long-term effects, which are also crucial in tES.

## Materials and methods

### 0.1 The Neocortical Computational Microscale Model (NeoCoMM)

NeoCoMM is a novel, physiologically-plausible, detailed model of a neocortical patch. It integrates the microcircuitry of the six cortical layers of the human neocortex (Figure 1) [35]. It takes into account the cyto-architecture of the neocortical column and includes the majority of cell types. These cells are divided into excitatory (Glutamatergic) and inhibitory (GABAergic) neurons. Excitatory neurons referred to as Principal Cells (PCs) consist of Tufted Pyramidal cells (TTPC), Untufted Pyramidal cells (UTPC), Inverted Pyramidal Cells (IPC), Bipolar Pyramidal Cells (BPC) and Spiny Stellate Cells (SSC). Inhibitory neurons referred to as interneurons (INs) include calcium-binding protein parvalbumin (PV+), neuropeptides somatostatin (SST+), vasoactive intestinal peptide (VIP+), and protein reelin (RLN+) expressing INs. PV+ expressing interneurons were further divided into Basket cells (BC) and Chandelier cells (ChC) (Figure 1.B) [35]. Each neuron cell was modeled using a reduced conductance-based model adapted to portray the firing dynamics of each cell type at different layers [35, 37]. Since it was shown that two compartments models can capture the impact of electric fields [9, 38, 39], PCs were portrayed by a three-compartment model including soma, dendrites, and axon initial segment. Each compartment includes a number of voltage-dependent ion channels, as shown in Figure 1.A (Left panel). Similarly, INs were represented by a mono-compartment model with a set of ion channels that portrayed the dynamics of each inhibitory type in the neocortical column [35, 37, 40]. The Hodgkin-Huxley formalism (Hodgkin and Huxley, 1952) was used to simulate the membrane potential of each compartment using the conservation of electric charge equation. A detailed description of single cell models used can be found in S1 Appendix. Cells were connected based on realistic connectivity rules and a predefined connectivity matrix. The connectivity diagram is portrayed by Figure 1.A (right panel). An example of a neocortical column simulated by NeoCoMM is given in Figure 1.C.

**Fig 1.**
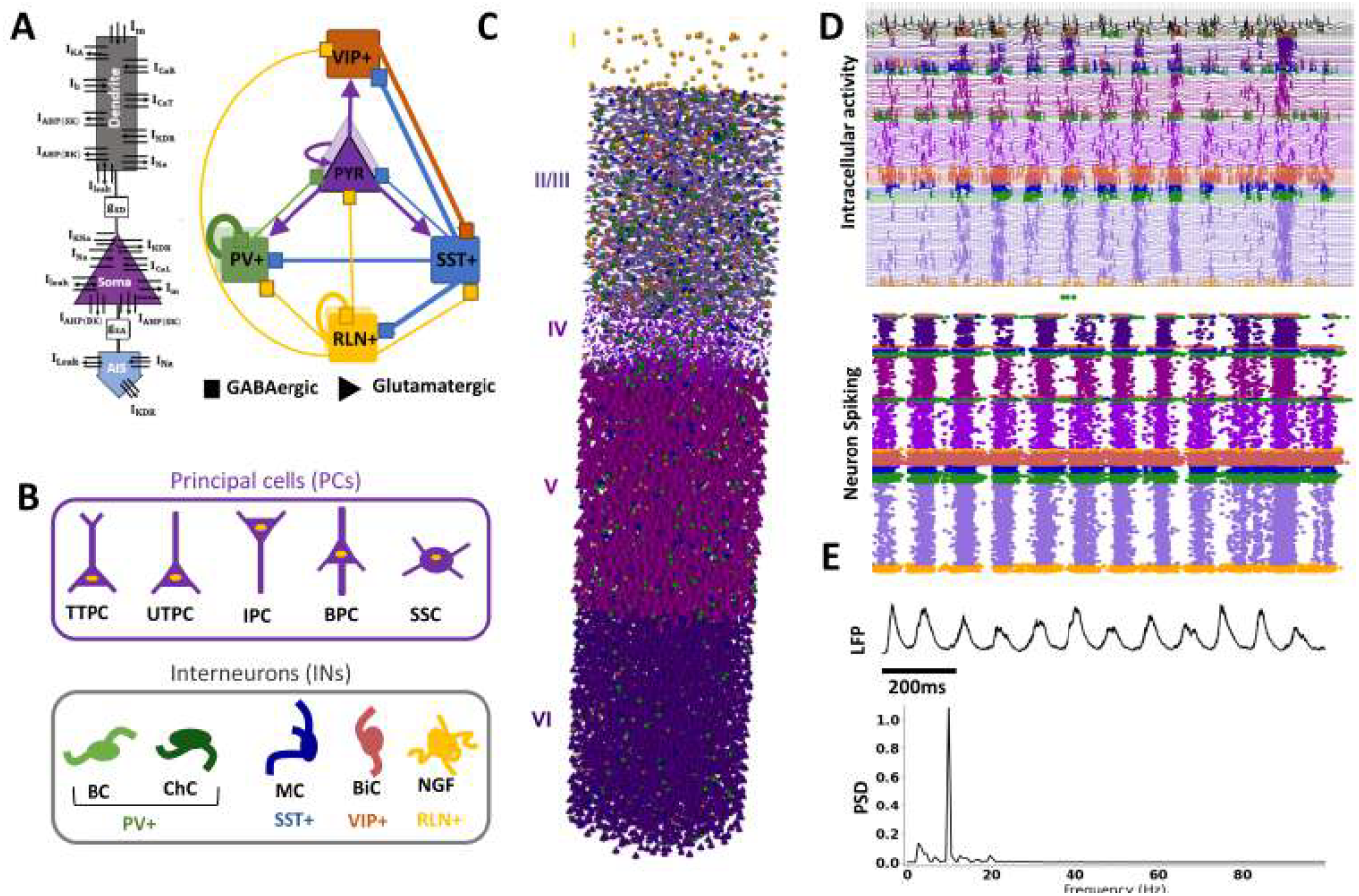
Network structure and simulation example with the NeoCoMM software. (A) The three-compartment neuron model (Soma, Dendrites, and Axon initial segment (AIS)) and the synaptic connectivity diagram. (B)The different cell types in the network: Tufted Pyramidal cells (TTPC), Untufted Pyramidal cells (UTPC), Inverted Pyramidal Cells (IPC), Bipolar Pyramidal Cells (BPC) and Spiny Stellate Cells (SSC), Basket cells (BC), Chandelier cells (ChC), Martinotti Cells (MC), Bipolar Cells (BiC) and Neurogliaform Cells (NGF). (C) A 3D example of the cortical column generated by NeoCoMM. (D) A simulation example of intracellular activity and neural spiking profile for an alpha rhythm (only 30% of cells’ activity are portrayed for visualization). (E) The simulated Local Field Potential (LFP) corresponding to the intracellular activity in (D).

The Local Field Potential (LFP) was estimated using a biophysical forward model of the volume conductor where the extracellular potential is computed and then integrated along the electrode surface (see [35] for details). Then, an electrode-tissue interface model [35, 36] was used to simulate the signal recorded by an intracerebral StereoElectroEnCephaloGraphic (SEEG) electrode typically used for the presurgical evaluation of patients. NeoCoMM is an open-source software package with a user-friendly interface available to download from https://github.com/MariamAlHarrach/Neocomm.V0.

### 0.2 Modeling cortical oscillations

Using the NeoCoMM Model framework, we were able to simulate realistic intrinsic cortical oscillations and epileptic events. This was done by adjusting the external input from the distant cortex (DC) and thalamus to the neocortical column along with the neurophysiological parameters of cell types and their synaptic connections. The parameter values corresponding to the different rhythms and events simulated are given in S2 Table.

#### 0.2.1 Endogenous rhythms

Without loss of generality, we simulated healthy continuous alpha-band and gamma-band rhythms that are among the frequency bands extensively studied in humans with scalp EEG (sEEG) to analyze the effect of tES on various healthy dynamics [41, 42]. This was done by adapting the stimulation input from the DC in the NeoCoMM framework. The input to PCs and INs in the cortical column consisted of a periodic volley of action potentials (APs) from PCs of the DC. The volley of APs was obtained by stimulating the PCs of the DC with injected currents to the soma with 66*µA/cm* ^2^ density and 30*ms* standard deviation for the alpha-band and 50*µA/cm* ^2^ density and 6*ms* standard deviation for the gamma-band simulations. To add heterogeneity and realism to simulations, the frequency of these volleys varied between 8 and 13*Hz* for alpha-band simulations and between 35 and 40*Hz* for gamma-band simulations. Physiological parameters (synaptic connections, conductances, and input stimulation) of the cortical network were determined based on the literature [35] and were adjusted using a sensitivity analysis and screening approach. These simulations respected the basal firing rate of individual cells as found in experimental studies and were one minute long each. All simulations are provided in the Supplementary Materials and can be run using the NeoCoMM software (https://github.com/MariamAlHarrach/Neocomm.V0).

#### 0.2.2 Epileptic events

In the case of the simulation of interictal epileptic events (IEEs), we refer to our previous work about the reconstruction and simulation of epileptic events using NeoCoMM [35]. We chose to simulate interictal spikes (IES), spikes and waves (SW), and high-frequency oscillations (HFOs). Based on [35], to simulate IEEs, two conditions are needed: (1) a hyperexcitable neural network and (2) a highly synchronized or quasi-synchronized stimulus (external input). For a more detailed description about the simulation of different epileptic events, please refer to our recent article [35] where we investigate the circuitry mechanisms responsible for the simulation of various interictal epileptiform events. The parameters adjusted in the NeoCOMM model to obtain the simulations used in this work are provided in Supplementary Table 2.

### 0.3 Modeling tES impact on network activity

To investigate the impact of tES on healthy and epileptic cortical activity, we modeled the contribution of tES based on a recent single-cell study [21] using detailed single-cell models within the NEURON environment [43] to compute polarization due to tACS with varying stimulation frequencies. These polarization length trends were then used to model tES in the NeoCOMM software with simplified cell models.

#### 0.3.1 Computing the single cell polarization lengths

In this section, we used the detailed NEURON models [43] to simulate extracellular stimulation via tACS. The simulation was implemented using the*extracellular* mechanism provided by the NEURON simulator [44]. The quasi-uniform approximation [45] was used to model electric field stimulation, assuming a uniform electric field aligned along the y-axis for all neuronal populations, including inhibitory neurons. This alignment corresponded to the somato-dendritic axis of pyramidal (PYR) cells. The extracellular potential was then calculated using Eq. 1.

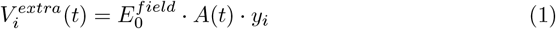

Where 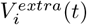 is the extracellular potential at the *i* ^*th*^ compartment at time *t, y* _*i*_ its coordinate, 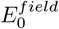 the unit electric field vector, and *A*(*t*) the stimulation waveform at time *t*.

The stimulation waveform was defined by *A*(*t*) =*a* · *sin*(2*πft*), where *a* is the electric field amplitude and *f*its frequency.

After initialization at the steady state without external inputs, two to four seconds simulations were performed for a stimulation frequency that spanned zero to 140 Hz (0.25 Hz step). The polarization length for each stimulation profile was defined as half the difference between the last maximum and minimum membrane potentials.

#### 0.3.2 Modeling tES in NeoCoMM

We defined a correspondence between the cell types included in NeoCoMM and those from the BBP [43] used to compute polarization length. The specific characteristics of these cells types, subtypes, types and morphologies are depicted in Supplementary Table 1. Morphologies are provided in S1 Fig Supplementary Figures 1 for PCs and 2 for INs. In brief, For PCs, we chose PYR cells for L23 and L4, TTPC for L5 and TPC, IPC and BPC for L6. For INs, PV+ cells were portrayed by LBC, SST+ by MC, VIP+ by BP and RLN+ by NGC.

In the case of PCs, we chose the morphology with the highest polarisation length for the soma compartment (Supplementary Table 1 and Supplementary Figure 1). Then, the polarization length for the dendrite compartment (simplified model) was obtained by averaging all apical and basal values across compartments. The polarization length for all PC compartments (dendrites + soma) is portrayed in Supplementary Figure 2. The frequency-dependent polarization lengths for all cell types are presented in Figure 3.

**Fig 2.**
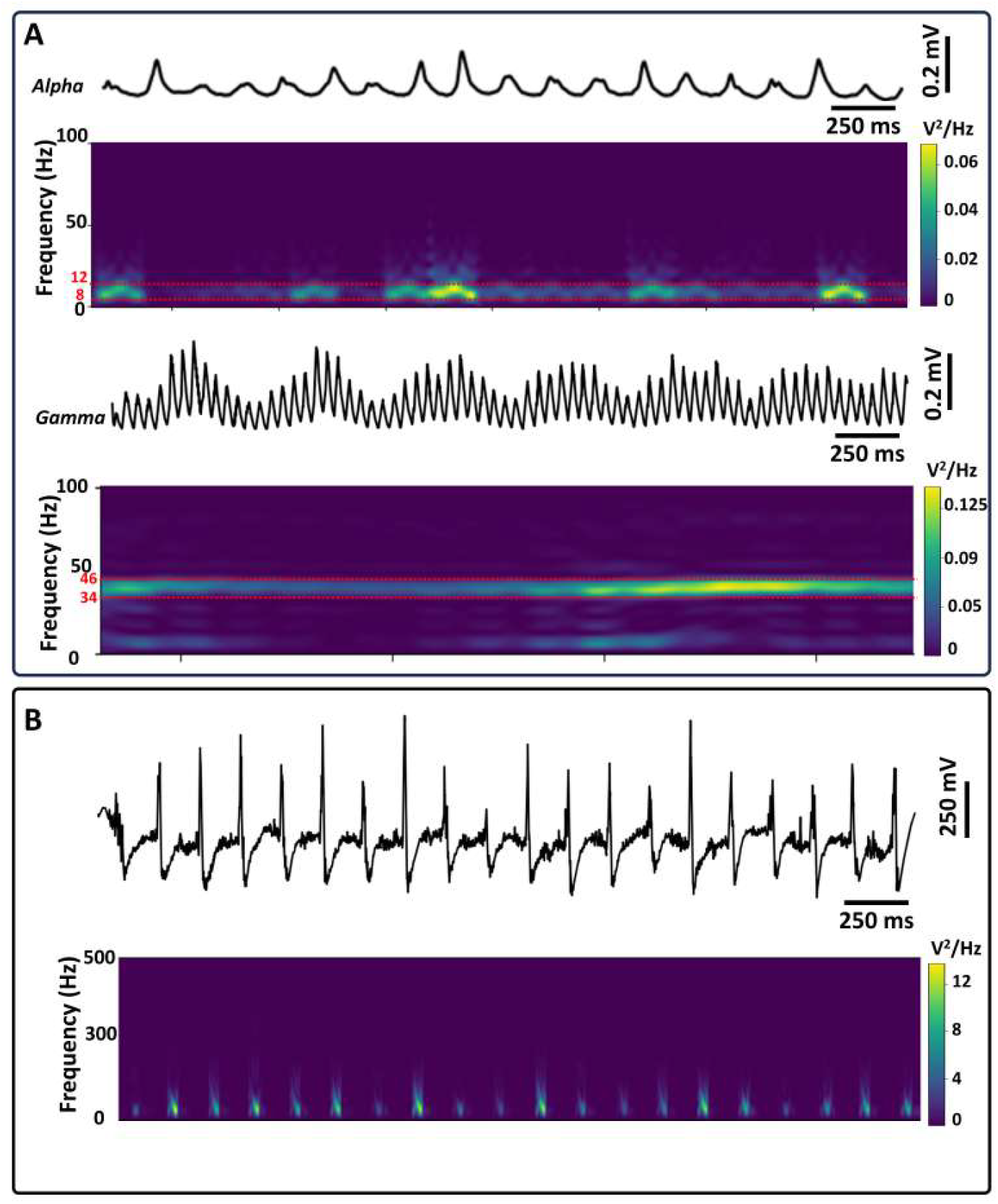
Example of two-minute simulations of healthy (A) and epileptic (B) rhythms using NeoCoMM. (A) LFP and corresponding power spectral density (PSD) of an Alpha (Top) and gamma (bottom) rhythm. (B) A LFP simulation of interictal epileptiform events (IEEs).

**Fig 3.**
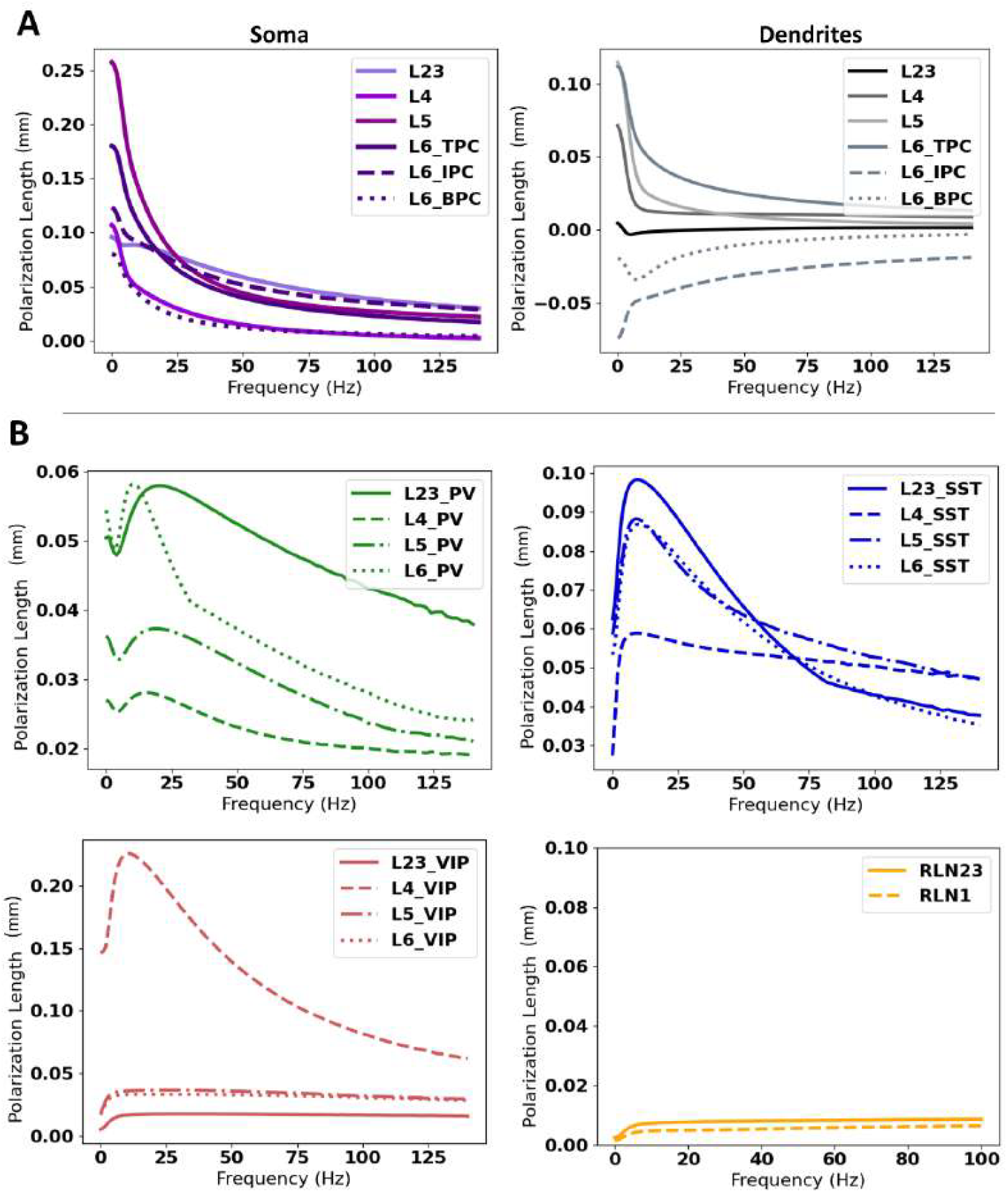
Frequency dependence of polarization length for all cell types. (A) Variation of the polarization length with tACS frequency for the soma (left panel) and the dendrites (right panel) of Principal Cells (PCs) of Layers L23, L4, L5, and L6. For L6 we investigated three subtypes of pyramidal cells: Tufted (TPC), Inverted (IPC), and Bipolar (BPC). (B) Variation of the polarization length with tACS frequency for the interneurons of the model (PV+, SST+, VIP+, and RLN+) at each layer (L23, L4, L5, and L6).

In the NeoCoMM model, the “lambda-E” formalism was used [46], which proposes that under weak and uniform electric fields, the membrane polarization across all compartments varies linearly with stimulation current amplitude. Consequently, the membrane polarization induced by the electric field at each compartment is represented as the dot product of the electric field vector and a length vector (polarization length) that characterizes the coupling constant, as shown in Eq 2. In the remainder of the manuscript, we will be referring to polarization length as *λ*.

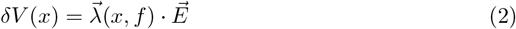

With the E-field vector, (x,f) the polarization length which is specific to each compartment and cell type portrayed by parameter *x* and *f* the stimulation frequency. For each cell type and layer, the corresponding polarization value was incorporated into the membrane potential, expressed as *V* ^*′*^(*x*) = *V* (*x*) + δ*V* (*x*) with *x* the parameter indicating the compartment and cell type.

### 0.4 Entrainment Analysis

Neural entrainment was quantified using the phase-locking value (PLV) [47], which quantifies the synchronization of spike timing relative to an ongoing signal. PLV is determined using Eq 3:

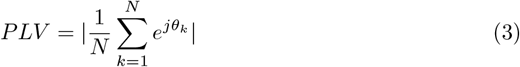

Here,*N* represents the total number of APs and*θ* _*k*_ denotes the phase of the tACS waveform at which the *k*-th spike occurs. Under the tACS condition, the PLV was computed relative to the tACS waveform. A *PLV* of 0 indicates a uniform distribution of spike timings across all phases (0 to 2*π*), while a value of 1 signifies complete synchronization to a specific phase of the tACS.

The Rayleigh test [48] was chosen to evaluate deviations from a circular uniform distribution. Polar plots were used to visually represent these quantitative values, as a preferred phase direction corresponds to higher bin counts in a specific angular direction.

To ensure that tACS impact portrayed by the PLV was significant compared to baseline, especially in the case of alpha rhythm where the network is already entrained to the 10 Hz frequency, We conducted a permutation test. Spike phases from stimulation and baseline were pooled and randomly reassigned into surrogate groups for 10,000 permutations to generate a null distribution of PLV difference between stimulation and baseline. PLV variation was considered as statistically significant for p ¡ 0.05.

### 0.5 Model simulation and data analysis

Simulations were performed using an expanded version of NeoCoMM . The software was run using a Dell workstation with Intel(R) Xeon(R) Gold (5220R CPU @ 2.20-4.00 GHz 2.19 GHz) processor using Pycharm Community Edition 2020.3.3 IDE with Python 3.10. No parallel processing was used. The simulations of different rhythms are available on the NeoCoMM GitHub public repository and can be loaded directly into the NeoCoMM software. As an example of computing time, for an alpha oscillation of 2s (1376 cells), running the simulation required approximately 15 minutes.

## 1 Results

The simulated cortical column networks used in the following analyses featured a total of 1,376 cells (see example in Figure 1.C). To explore oscillatory dynamics, we tested various network configurations capable of generating endogenous alpha and gamma rhythms across a range of frequencies within their respective bands (examples shown in Figure 2). For epileptiform activity, we modeled a series of epileptic events emerging from a highly excitable network. Each simulation ran for 4 minutes: a 2-minute baseline period without tACS, followed by 2 minutes with tACS. We examined both the extracellular field potentials, as measured by a virtual electrode designed to mimic SEEG recordings (see Section 2.2), and the intracellular activity across different neuronal cell types. Results are detailed in the following subsections.

### 1.1 tACS Impact on Physiological Cortical Oscillations

We quantified the neural entrainment of PCs and INs across stimulation frequencies of 5, 10, 20, 30, 40, 80, and 120*Hz*, and electric field intensities of 0.3, 0.5, 1, and 3*mV/mm*. For PCs, we grouped together those with similar electrophysiological properties: specifically, layer 2/3 and layer 4 PCs (PC23 and PC4), as well as layer 5 and layer 6 tufted PCs (PC5 and TPC6). For clarity, we present results for only one representative from each group. However, for layer 6, we also report results for inverted (IPC) and bipolar (BPC) pyramidal cells, since their morphologies differ in terms of directionality from the other PC types.

To investigate the effect of alpha spectral peak frequency on tACS-induced entrainment, we simulated alpha oscillations at 8, 9, 10, 11, and 12*Hz*. The baseline power spectral densities for each condition are shown in Supplementary Figure 4 (left). For the simulation at 10*Hz* peak frequency, PCs had a mean firing rate of *±* 7.55 2.43 Hz. Figure 4.A presents the variation of phase-locking value (PLV) across stimulation frequencies and intensities for pyramidal cells (PCs). In the absence of stimulation, the baseline simulation already exhibited a high PLV at 10*Hz*, reflecting the presence of a strong endogenous 10*Hz* peak. Under tACS, the LFP showed entrainment at this frequency, beginning at a stimulation intensity of 0.3 *mV/mm*. We also observed weaker entrainment at 20 *Hz* and 30 *Hz*. This was observed for all the PC layers. However, permutation testing demonstrated that PLV values were significantly elevated relative to baseline only when the applied tACS amplitude exceeded 1,mV*/*mm . At a stimulation frequency of 10,Hz, this effect was restricted to pyramidal cells located in layers 5 and 6, whereas at 20 and 30,Hz, significant PLV increases were observed across pyramidal cells in all cortical layers (Figure 4). The corresponding preferred phase (PPh) distributions are shown in Figure 4.B. Notably, across stimulation frequencies, the PPh varied but tended to converge toward a consistent phase as stimulation intensity increased, suggesting phase alignment driven by tACS. Polar plots in Figure 4.C further illustrate strong neural entrainment at 10*Hz* with slightly different PPh values across the different PC subtypes, confirmed by a highly significant Rayleigh test (p*<*0.0001). Supplementary Figure 5 shows polar plots of the PPh at baseline and during 1,mV*/*mm tACS at different stimulation frequencies. The plots demonstrate clear frequency-dependent shifts in the PPh, indicating that tACS entrains neuronal activity with distinct phase preferences depending on the stimulation frequency.

**Fig 4.**
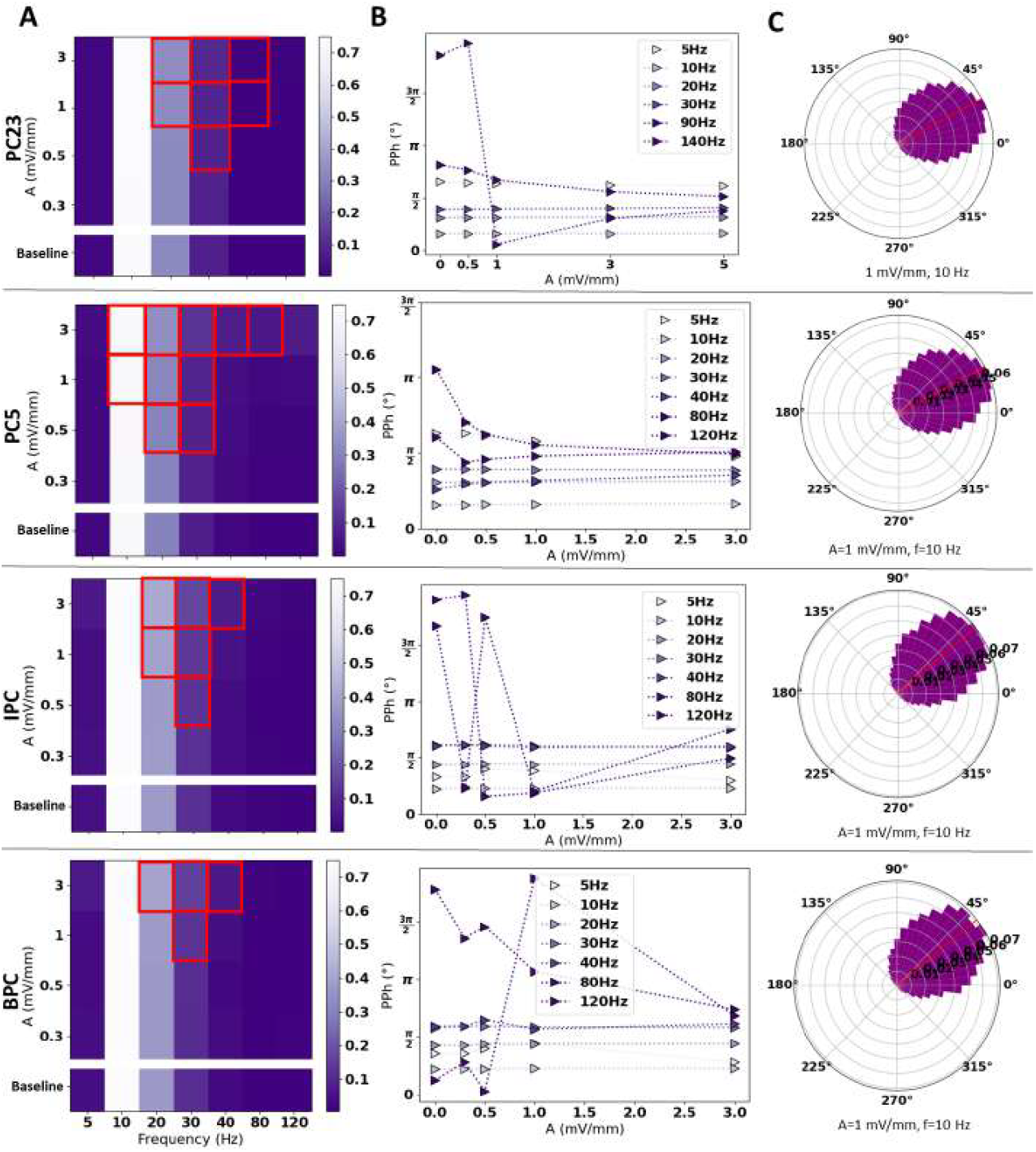
Impact of tACS on the network activity of Principal Cells (PC) of layer 23 (PC23), layer 5 (PC5), Inverted PCs of layer 6 (IPC) and Bipolar PCs of layer 6 (BPC) on 10 Hz alpha rhythm. (A) Entrainment maps of PCs to tACS with varying intensity and frequency. Red rectangles indicate conditions in which the phase-locking value (PLV) during tACS was significantly greater than baseline (permutation test, p ¡ 0.05). (B) Preferred Phase (PPh) variation with tACS parameters. (C) An example of entrainment of PC’s action potentials for 1mV/mm, 10Hz tACS.

**Fig 5.**
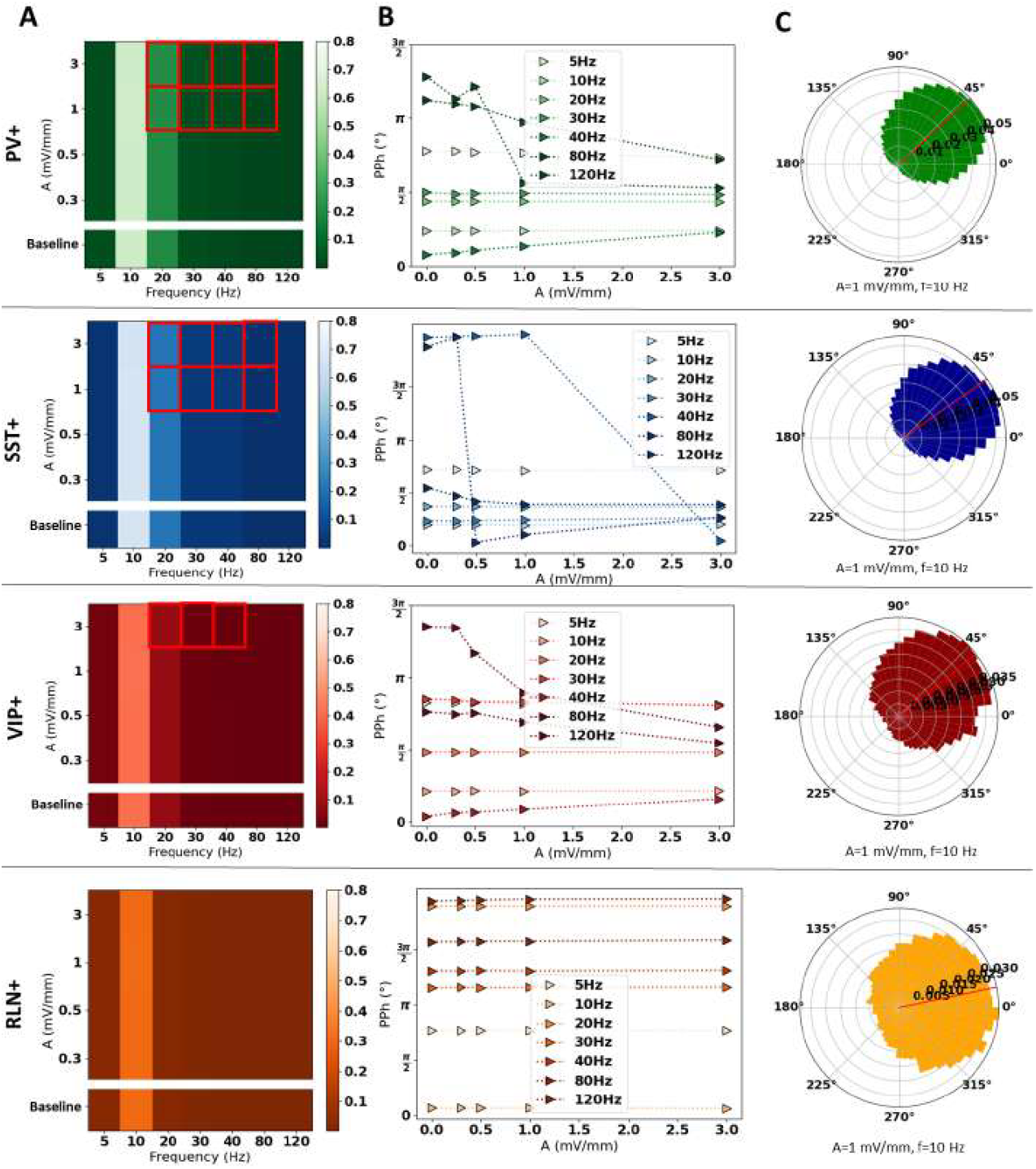
Impact of tACS on the network activity of Interneurons (INs) of layer 23. (A) Entrainment maps of PCs to tACS with varying intensity and frequency. Red rectangles indicate conditions in which the phase-locking value (PLV) during tACS was significantly greater than baseline (permutation test, p ¡ 0.05). (B) The Preferred Phase (PPh) variation with tACS parameters. (C) An example of entrainment of PC’s action potentials for 1 mV/mm, 10 Hz tACS. PV+: parvalbumin expressing INs, SST+: somatostatin expressing INs, VIP+: vasoactive intestinal peptide expressing INs, RLN+: reelin expressing INs.

Figure 5 presents tACS entrainment results for the same alpha-band simulations, focusing on the four types of interneurons (INs) modeled in the network. All IN types exhibited entrainment at 10 Hz, with SST^+^-expressing INs showing the highest PLV values and RLN^+^-expressing INs the lowest (Figure5.A). The preferred phase (PPh) varied depending on both the interneuron type and stimulation frequency. At higher stimulation frequencies, the PPh tended to converge towards a distinct value as the stimulation intensity increased, as illustrated in Figure 5.B.

Polar plots indicate stronger phase locking and entrainment in PV^+^ and SST^+^ INs compared to VIP^+^ and RLN^+^ INs. Baseline mean firing rates were 43.25 ± 15.36, 50.09 ± 6.87, 44.06 ± 6.26, and 67.43 ± 13.73 *Hz* for PV^+^, SST^+^, VIP^+^, and RLN^+^ INs, respectively. While these rates are considerably higher than those of PCs, the strongest entrainment to 10*Hz* tACS was still observed at this frequency. Similary to PCs, the permutation tests on the PLV only revealed significant variation compared to the baseline for tACS amplitude above 0.5*mV/mm* for PV ^+^ and SST^+^ INs and above 0.5 *mV/mm* for VIP^+^. For RLN ^+^ the PLV values were not significantly different from the baseline. Despite this, the PPh and phase polar plots clearly show a phase entrainment at 10, 20*Hz*.

We performed the same analyses for alpha oscillations with different peak frequencies within the alpha band. For simulations with a 8 *Hz* peak alpha frequency, the mean firing rate (FR) of pyramidal cells (PCs) was 6.40 ± 2.07 Hz. These cells exhibited reduced sensitivity to tACS, as indicated by low PLV values and weaker overall entrainment. PLV maps for PCs under these conditions are shown in Supplementary Figure S6, alongside the PPh distributions and polar plots for 10 *Hz* tACS at 1 *mV/mm*. While the overall entrainment was lower, PCs still showed a gradual convergence towards a specific preferred phase as stimulation intensity increased. However, the Rayleigh test for 10 *Hz* stimulation at 1*mV/mm* was less significant compared to the 10*Hz* peak simulations, and PPh remained more variable across PC subtypes.

In the case of gamma oscillations, we applied the same analysis as for the alpha band. Power spectral densities (PSDs) of the five different gamma rhythms used in this study are presented in Supplementary Figure S7 (right), covering frequencies from 36 to 50 Hz.

Figure 6 illustrates the effect of tACS on gamma oscillations with peak frequencies at 40 Hz and 50 Hz, in terms of phase locking and entrainment for PC cells of layer 5 (PC5). Comparing the PLV maps for these two gamma frequencies, we observed strong entrainment when tACS frequency matched endogenous gamma rhythm—specifically, at 40 Hz for the 40 Hz gamma case (Figure 6.B). In contrast, for 50 Hz gamma oscillations, PLV values were generally low, with the highest values occurring at 10 and 20 Hz tACS, and only at stimulation intensities above 0.3*mV/mm*. Permutation test revealed that the PLV was significantly different from the baseline value when the tACS amplitude was above 0.5*mV/mm*.

**Fig 6.**
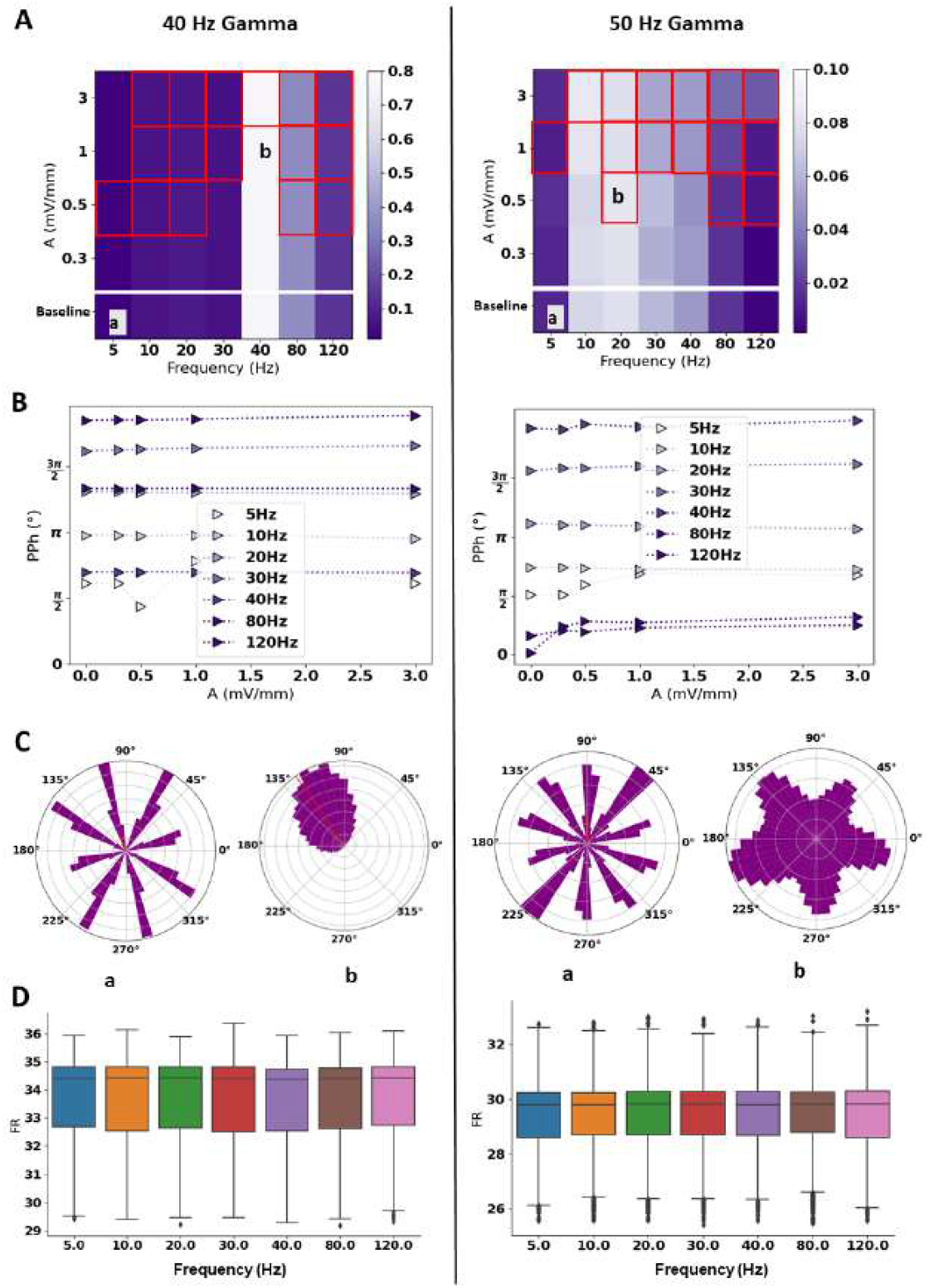
Impact of tACS on layer 5 principal cells (PCs) in a network gamma activity at two frequencies, 40 Hz (left) and 50 Hz (right). (A) Entrainment maps of layer 5 PCs to tACS with varying intensity and frequency. Red rectangles indicate conditions in which the phase-locking value (PLV) during tACS was significantly greater than baseline (permutation test, p ¡ 0.05). (B) The Preferred Phase (PPh) variation with tACS parameters. (C) Comparison of the entrainment of PC’s action potentials at baseline versus max entrainment for 40 Hz (left) and 50 Hz (right) gamma activity. (D) Box plots of the firing rate (FR) variation with tACS frequency. PV+: parvalbumin-expressing INs, SST+: somatostatin-expressing INs, VIP+: vasoactive intestinal peptide-expressing INs, RLN+: reelin-expressing INs.

PPh in both gamma rhythm simulations depended on tACS frequency, as shown in Figure 6.B. In the 40 Hz gamma case, the PPh remained relatively stable across frequencies. In the 50 Hz gamma case, small shifts in PPh were observed at lower tACS intensities.

Phase polar plots in Figure 6.C further highlight the strong entrainment when stimulation matched the endogenous frequency (40 Hz), relative to the baseline. This was confirmed by the Rayleigh test (P *<*0.001). In the case of tACS applied at different frequencies as compared to the endogenous frequency, as is the case for 50 Hz gamma oscillations, entrainment was less significant, and the phase polar plot for the tACS configurations with the highest PLV (tACS at 20 Hz, 0.5 mV/mm) confirmed this result.

Regarding the firing rate, we did not observe any significant changes between different tACS frequencies or intensities. Figure 6.D presents the firing rate boxplots for PC5 cells under 1 *mV/mm* tACS across the seven tested frequencies. It is worth noting that, in the 50 Hz gamma simulations, a subgroup of PC5 cells fired at around 25 Hz. This could help explain the relatively higher PLV values observed with 20 Hz tACS in Figure 6.A. Similar results were observed for the remaining principal cells including P23, PC6, IPC and BPC.

### 1.2 tACS Impact on Epileptic Events

In this section, we investigated the influence of transcranial alternating current stimulation (tACS) parameters on epileptic activity. To this end, we simulated two minutes of epileptic activity by making the network hyperexcitable and delivering external inputs from both the distant cortex and thalamus. The parameters used for these simulations are detailed in Supplementary Table 2.

To quantify the impact of tACS on epileptic discharges, we extracted three features from spike-like events: (i) percentage variation in the number of epileptic events, (ii) spike amplitude, and (iii) full width at half maximum (FWHM). The latter two features were computed using the *find_peaks* function from the *scipy* library (Python 3.12.0).

Figure 7 illustrates the effects of tACS on interictal epileptic events (EEs) across different stimulation intensities and frequencies.

**Fig 7.**
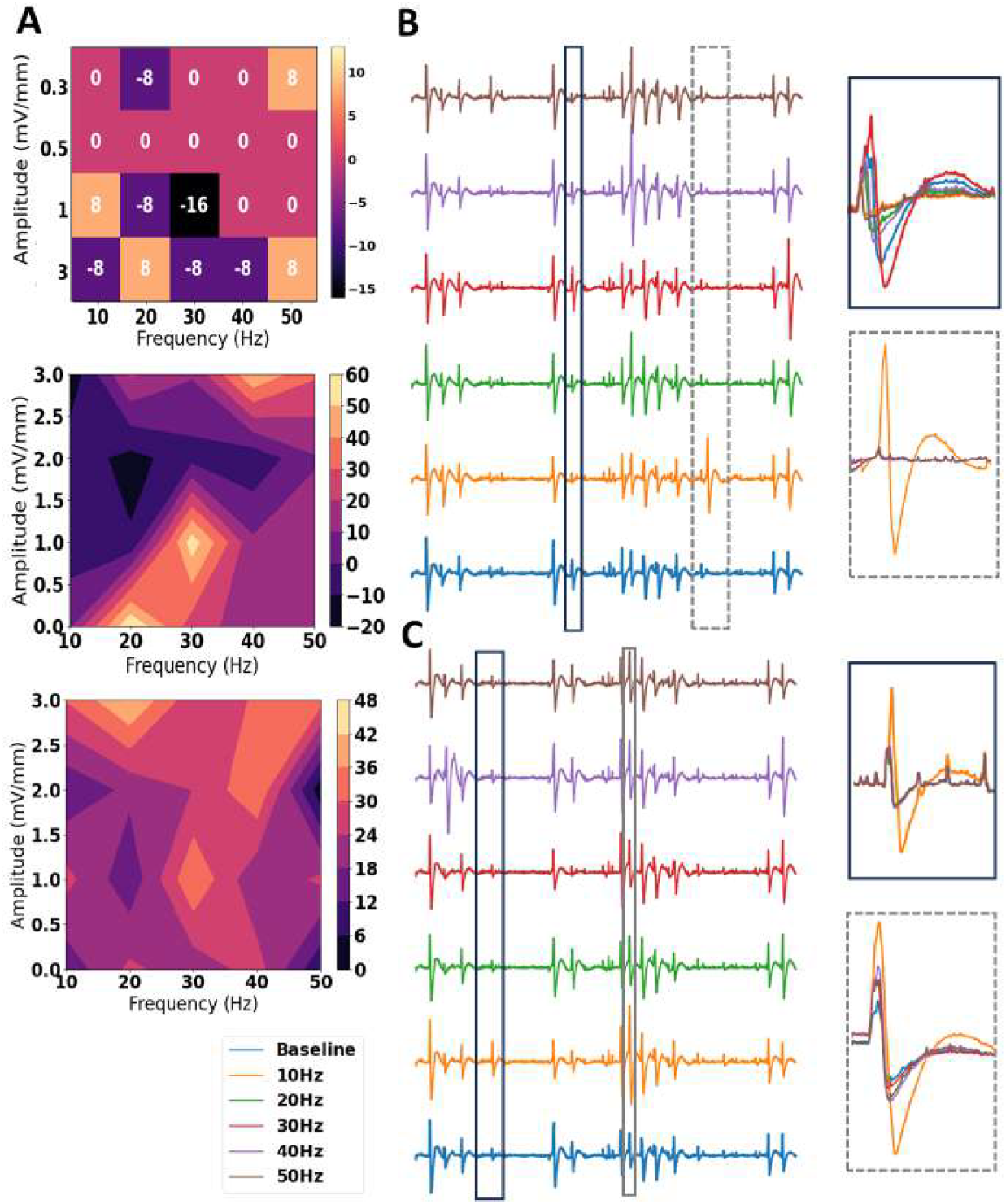
Impact of tACS on Epileptic activity. (A) Variation of the percentage (Top), mean maximum peak amplitude (middle), and Full Width at Half Amplitude (FWHA) (bottom) of interictal epileptic events (EEs) with frequency and amplitude of tACS. (B) Example of local field potential (LFP) variation with frequency at 1 mV/mm and (C) 0.5 mV/mm (bottom).

As shown in Figure 7.A, there was no consistent trend in the number of EEs across all stimulation conditions. However, at 3*mV*/mm tACS, we observed a noticeable reduction in the number of EEs for stimulation frequencies of 10, 30 and 40 *Hz*. Lower stimulation intensities had a weaker overall effect on EEs occurrences.

Regarding spike amplitude (Figure 7A), we observed the largest increase at 30 Hz, which was accompanied by an increase in FWHM, suggesting broader and more pronounced spike events.

Figures 7.B and C present representative signal traces for tACS at 1 mV/mm and 0.5 mV/mm, respectively. At 1*mV/mm*, we observed an increased number of discharges at 10 *Hz* (first zoomed segment); however, spike amplitude decreased at the same frequency (second zoomed segment). In the 0.5 *mV/mm* condition, the number of events remained largely unchanged, but spike amplitude exhibited a global increase, most notably at 10 *Hz* in some individual spikes (Figure 7C). These results suggest that the impact of tACS on epileptic discharges is highly dependent on both stimulation parameters and specific characteristics of the events.

### 1.3 tDCS Impact on Epileptic Events

We also performed a comparable analysis for transcranial direct current stimulation (tDCS), examining both cathodal (ctDCS) and anodal (atDCS) stimulation at an intensity of 0.5 mV/mm. Results are shown in Figure8, based on the same simulation setup described in the previous subsection.

As expected, ctDCS resulted in a reduction of spike amplitude, while atDCS led to an increase. However, these opposing effects were not symmetrical in magnitude. Notably, ctDCS also led to a decrease in the number of epileptic events, a change that was not observed with atDCS. This asymmetry suggests that the underlying mechanisms by which ctDCS and atDCS modulate epileptic activity may differ in sensitivity or effectiveness. To illustrate these effects, we selected two representative epileptic spike events from the signal traces shown in Figure 8.A. In the first example (purple), we observed an overall increase in both the amplitude and duration of the event under atDCS, while ctDCS produced no notable change. The corresponding raster plot of PC5 firing activity (Figure 8.B) shows a clear increase and temporal shift in firing rate under atDCS, whereas ctDCS resulted in minimal alteration.

**Fig 8.**
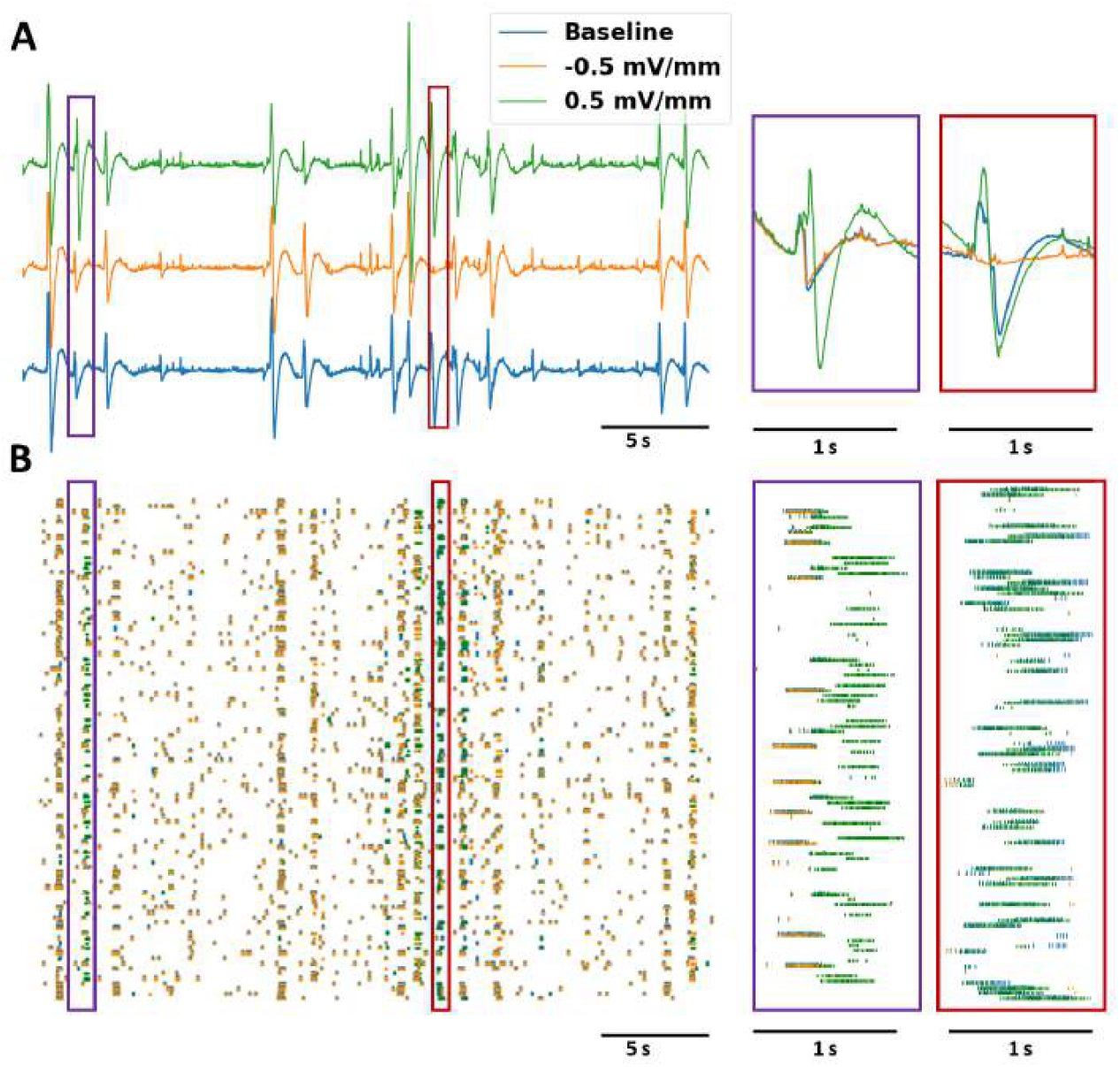
Example of cathodal versus anodal tDCS impact on epileptic spike-like activity in the neocortex. (A) Comparison of 0.5 and -0.5 mV/mm tDCS effect on simulated epileptic discharges. (B) Raster plot for layer 5 Principal cells’ (PC5) action potentials (APs).

In the second example (red), ctDCS effectively abolished the epileptic spike, as evidenced by the absence of action potentials during the corresponding segment in the raster plot (Figure 8B). In contrast, atDCS increased the amplitude of the spike without significantly affecting its duration. This was reflected in the raster plot, where a higher degree of synchronization among PC5 neurons was observed under atDCS (red zoomed rectangle in Figure 8.B).

## Discussion and conclusions

In this study, we extended a recently developed neocortical computational microscale model, NeoCoMM, to study the impact of tES on the activity of a network of cortical cells in a multilayered cortical column. Specifically, we examined how tDCS and tACS modulate extracellular and intracellular activity in both healthy and epileptic network configurations. Furthermore, we systematically explored how variations in stimulation frequency and intensity influence cortical network activity.

Compared to the recent study by Zhao et al. [9], we did not observe a de-synchronization at lower stimulation intensities for the alpha oscillation. Entrainment was consistent even for low stimulation magnitudes (0.3 *mV/mm*) (Figures 4, 5,6). In their study, Zhao et al. observed a de-synchronization relative to baseline for electric field magnitudes below 0.6 *mV/mm* in the case of 10 Hz alpha oscillations, but only below 0.2*mV/mm* in the case of low beta [9]. This difference might be explained by the added complexity of the model, which encompasses more inhibitory neuron types and layer-specific cells and connectivity as compared to the two-cell network used in [9]. Moreover, to verify that the observed impact of tACS in our study on network entrainment was not due to baseline oscillation, we conducted a permutation test on firing phase values. This test revealed that PLV values were significantly different from the baseline for tACS intensities for specific amplitudes and frequencies of tACS. Nevertheless the PPh did reveal that the network synchronisation is directly dependent on the tACS frequency.

When endogenous frequency differed from the target stimulation frequency, entrainment was less significant. This was the case for the 8 Hz alpha stimulation for a 10 Hz tACS (Supplementary Figure 4), and the 50 Hz gamma stimulation for the 40 Hz tACS (Figure 6). This resonance-like phenomenon was also observed in the Zhao et al. study [9], highlighting the possibility that LFP power peak frequency determines the tACS frequency that causes the highest entrainment, and not the individual firing rate of cell types as demonstrated in single cell analyses. This could also explain why some cells in a network are more entrained, which is well portrayed by Figure 6. In this case, stimulation at 40 Hz clearly entrained PC5 cells in the case of 40 Hz gamma (peak power at 40 Hz as portrayed by Supplementary Figure 4) and especially in the baseline versus tACS at 1*mV/mm*, 40 Hz phase polar plot. However, entrainment was not significant for 50 Hz gamma, and was highest around 10-20 Hz which does not relate to the firing rate of individual PC5 cells. However, since we did not test tACS at 50 Hz we cannot conclude that this exact stimulation frequency would not produce strong entrainment. Future analysis with increased tACS intensity/frequency parameter resolutions should be tested in the future to confirm this hypothesis.

In terms of cell type sensitivity to stimulation, we did not observe high sensitivity for a particular cell layer or type. We did find subtle differences between the preferred entrainment phase for different stimulation frequencies. Also, PLV was highest for PV+ and SST+ interneurons compared to other INs and even PCs (Figures 4 and 5). For layer 6, PCs of the IPC and BPC types showed high phase variation for low stimulation magnitudes that only converged at 3*mV/mm*. This can be explained by the polarization length that are maximal around 10 Hz for PV+ and SST+ INs as compared to other cell types (Figure 3). This further validates the hypothesis that cell-class-specific tES entrainment studies on single cells do not necessarily translate to network activity case, such as in [6].

In addition, all these different stimulation scenarios did not result in a change in the firing rate of cells with increased intensity or resonance value frequencies. This is in accordance with recent studies on electric field entrainment of network activity and single cells mainly in the cortex at subthreshold stimulation [3, 6, 9, 21, 49].

Since the effects of tDCS on physiological alpha and gamma rhythms have already been well established in the literature [50–52], we did not include a detailed analysis of these effects in the main text. Nevertheless, Supplementary Figure 6 presents the results of tDCS applied at different field intensities, showing no observable changes in signal morphology, neuronal firing rates, or power spectral density (PSD). However, it is important to note that a recent in vivo study by Farahani et al. [53] found that rats exposed to tDCS at clinically relevant field strengths experienced notable changes in hippocampus cell firing rates. This result draws attention to the possible sensitivity of particular brain areas or states to direct current stimulation, which the cortical context might not adequately represent.

In the case of an epileptic network, we studied the intensity and frequency specific impact of tACS on a simulation with spike-like Epileptic Events (EEs) (Figure 7). Results revealed inconsistent impact on epileptic events with increased stimulation intensity. At the highest intensities, tACS reduced the mean number of EEs at 10 and 50 Hz. However the mean peak amplitude and FWHA were not reliably impacted. This suggests that tACS impact depends on the event and the underlying network and cell dynamics responsible for this epileptic event. This is more discernible in the zoomed events portrayed by Figure 7.B. In the case of 10 Hz tACS stimulation, an additional EE is produced compared to the baseline simulation. However, in the same simulation, the last EE, presented a decrease in its width and amplitude. Then, for a slight increase in tACS intensity, the network presents a totally different response to the same frequencies; confirming the concerns presented in some studies about the use of tACS in epilepsy [54, 55]. It also supports the hypothesis that we formulated previously regarding the important role of network activity on the choice of tACS parameters and its impact on individual cells in a network setting. This is especially important to consider when using results from studies in a specific network activity or configuration as in [56].

We also tested the impact of both cathodal and anodal tDCS on the same signal, and found as expected a non-symmetrical result where ctDCS decreased the number, amplitude and width of EEs, and atDCS increased the already existing EEs amplitude and duration (8). The effect is explained by decreasing the firing rate of excitatory cells, and in some cases stopping their bursting activity for ctDCS. These results are aligned with many studies about the efficacy of tDCS in reducing seizures and interictal epileptic events [16, 57].

In conclusion, our study presents a new open-source model that can be used to test and optimize tES parameters in a specific simulation scenario for a cortical network of cells. Here, it was used to study the effects of tACS on physiological alpha and gamma rhythms as well as epileptic activity. The main results suggest that, in a physiological (healthy) network tACS, parameter optimization depends on network activity, and more specifically on extracellular activity (LFP). Therefore, the cell- and frequency-specific tACS recommendation should be tailored to network settings and activity. In the case of a hyperexcitable network, tACS does not provide conclusive results, which further validates the specificity hypothesis mentioned above. This study advances our understanding of the underlying mechanisms of tES in general and tACS in particular, where computational modeling is considered crucial for tACS parameter and protocol design and optimization [3]. Furthermore, we quantified how the electric field interacts with a network of cells and can produce different results depending on the activity produced by the network and individual cells. These insights will be instrumental for the optimization of tES for specific therapeutic neuromodulation therapies [58, 59].

## S1 Appendix

A spatially reduced, three-compartment model was employed to represent Principal Cells (PCs), whereas interneurons (INs) described using a single-compartment model. The three-compartment PC model followed our previous framework [35], and the single-compartment IN model was adapted from Hajos et al. (2004). These reduced representations provide an effective balance between physiological fidelity and computational efficiency. In the PC cells model, the three compartments correspond to the soma, dendrites and Axon Initial Segment (AIS). These compartements are coupled via two conductances (Figure 1.A). The membrane potential dynamics in each compartment were simulated using the Hodgkin–Huxley formalism [60] and governed by the standard charge-conservation equation below:

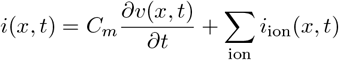

with *i*(*x, t*), *v*(*x, t*) and *i*_*ion*_(*x, t*) are the injected input current, membrane potential, and ionic currents of compartment *x*, respectively. The ionic currents were compartment- and cell-type–specific and were defined as

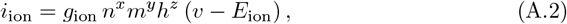

where *g* _ion_ denotes the maximal ionic conductance, *E* _ion_ the reversal potential, and *m, n*, and *h* are gating variables representing the probability that a channel is open at a given time. The exponents *x, y*, and *z* are positive integers corresponding to the number of identical gating subunits. Variables *m* and *n* describe channel activation, whereas *h* represents inactivation. The temporal evolution of the activation and inactivation variables for each channel is given by:

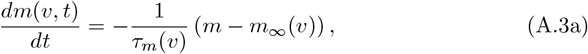

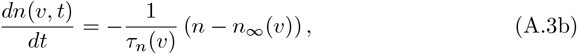

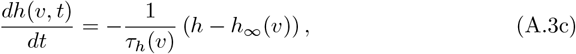

where the steady-state activation and inactivation functions and their time constants are given by:

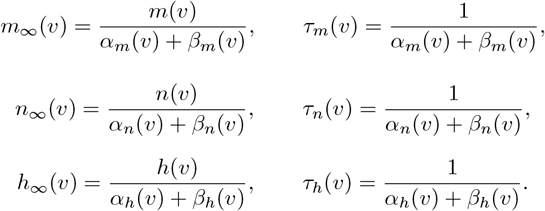

Ionic channel parameter values (mainly conductances) were adjusted to reproduce the firing rate obtained through electrophysiological recordings. A detailed description of ionic current equations and parameter values can be found in [35].

**S1 Fig.**
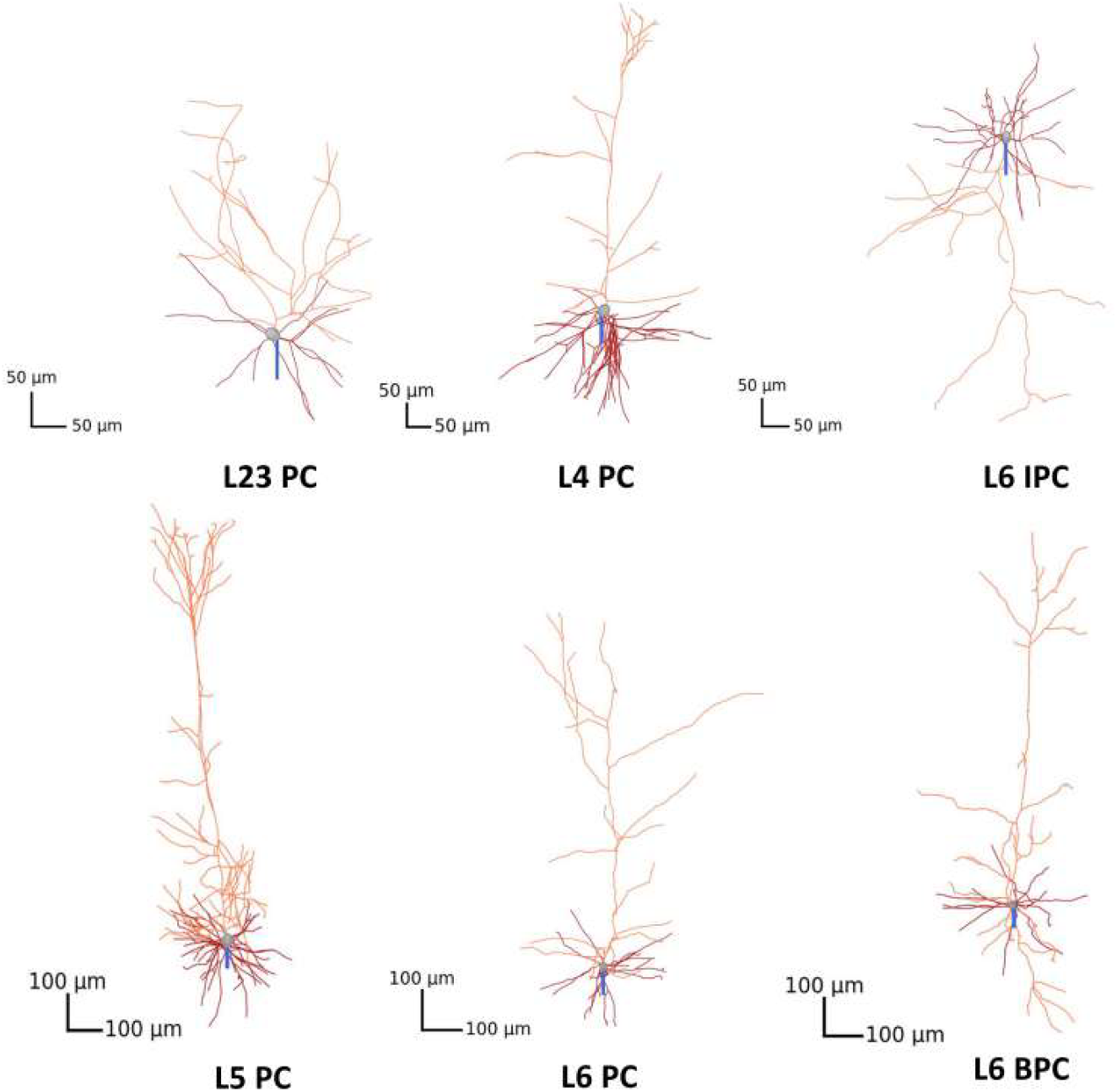
Morphologies of the Principal Cells (PCs) used to compute the polarization length *λ* variation with tES frequency. These PC types are as follow: pyramidal (PYR) cells for L23 and L4, Tufted Pyramidal cells (TTPC) for L5, and Tufted Pyramidal cells (TPC), Inverted Pyramidal cells (IPC) and Bipolar Pyramidal cells (BPC) for L6. The corresponding morphology name and e-type in [43] are given in S1 Table.

**S2 Fig.**
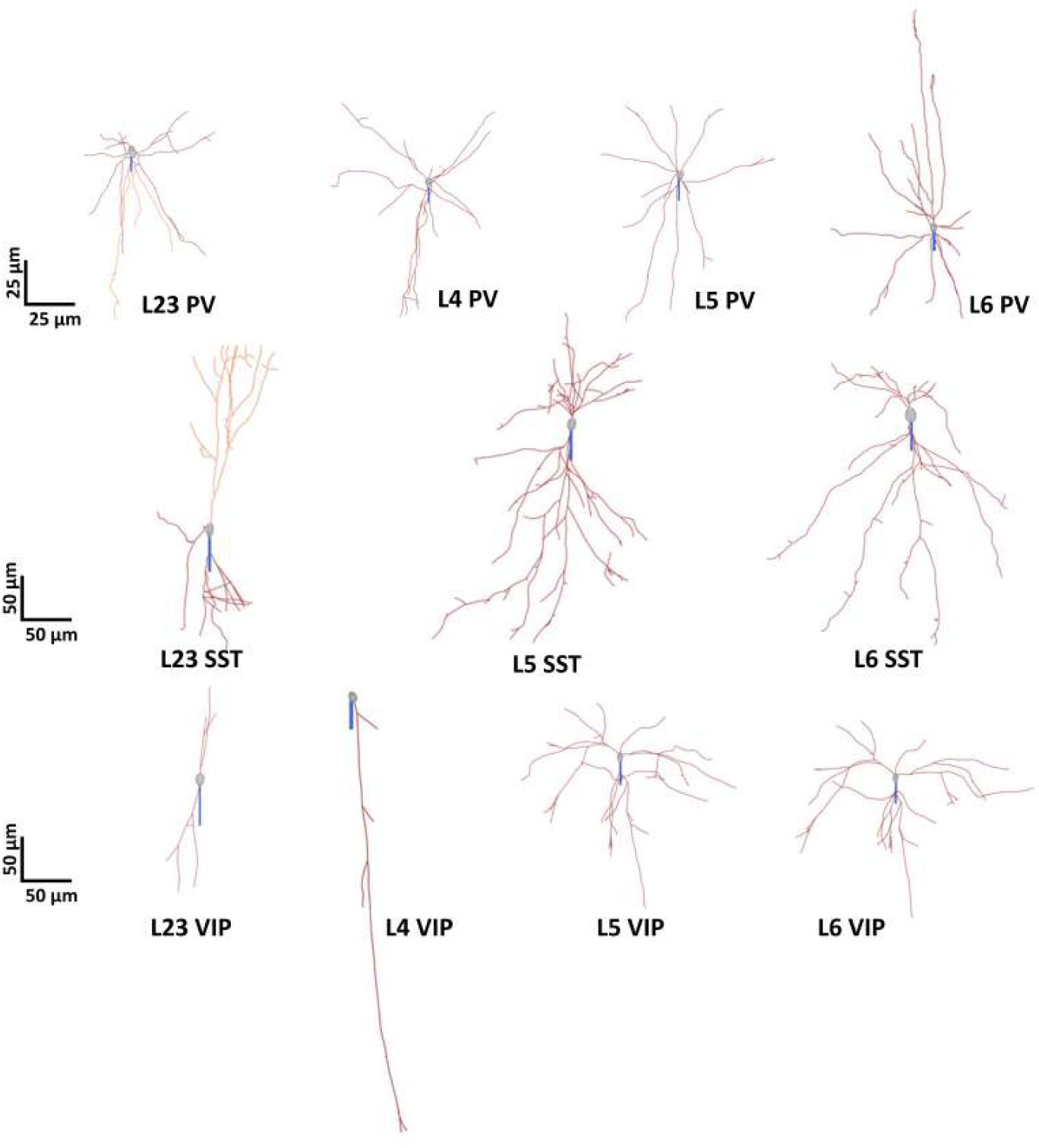
Morphologies of the Inhibitory Cells (INs) used to compute the polarization length *λ* variation with tES frequency. The corresponding morphology name and e-type in [43] are given in S1 Table.

**S3 Fig.**
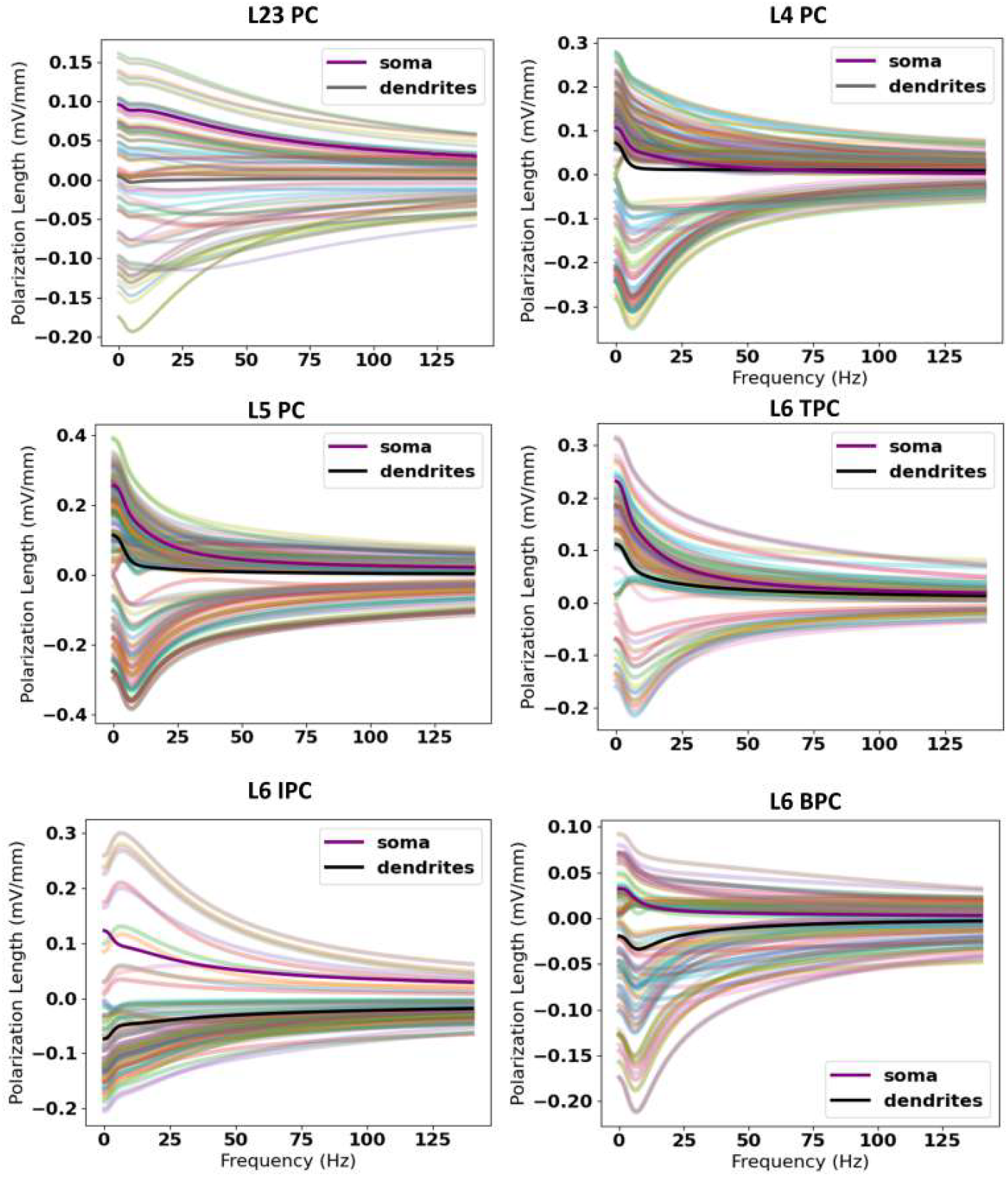
Frequency dependence of polarization length for all cell types. (A) Variation of the polarization length with tACS frequency for the soma (left panel) and the dendrites (right panel) of Principal Cells (PCs) of Layers L23, L4, L5, and L6. For L6 we investigated three subtypes of pyramidal cells: Tufted (TPC), Inverted (IPC), and Bipolar (BPC). (B) Variation of the polarization length with tACS frequency for the interneurons of the model (PV+, SST+, VIP+, and RLN+) at each layer (L23, L4, L5, and L6).

**S4 Fig.**
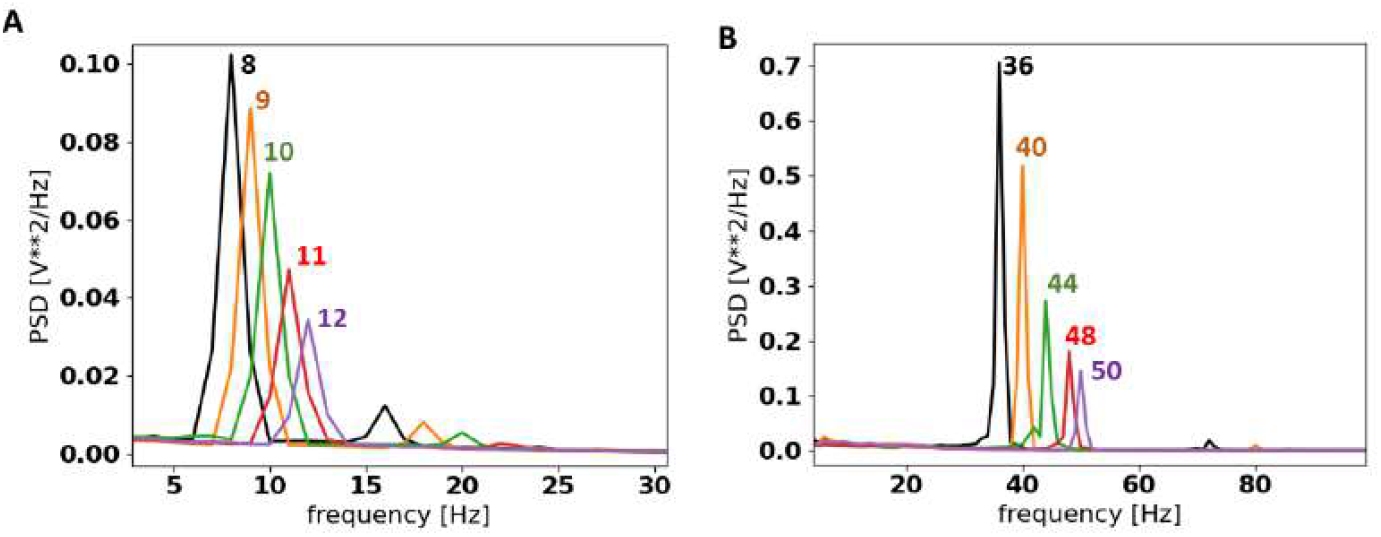
Power spectral density of the alpha (A) and gamma (B) rhythm signals simulated in the study.

**S5 Fig.**
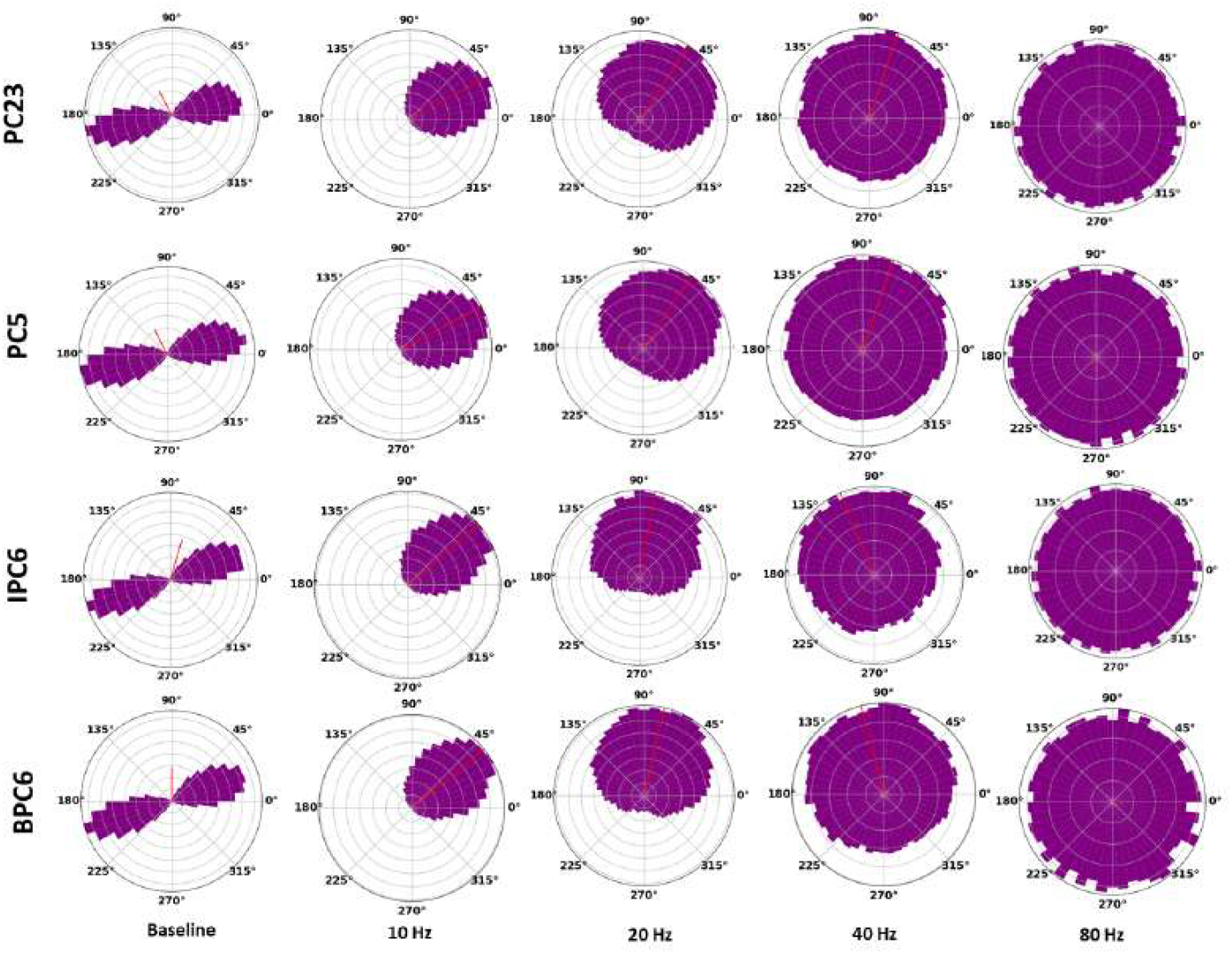
Polar plots of the Prefered Phase (PPh) of Principal Cells (PC) of layer 23 (PC23), layer 5 (PC5), Inverted PCs of layer 6 (IPC) and Bipolar PCs of layer 6 (BPC) at Baseline compared to 10, 20, 30, 40 and 80*Hz* tACS at 1*mV/mm*.

**S6 Fig.**
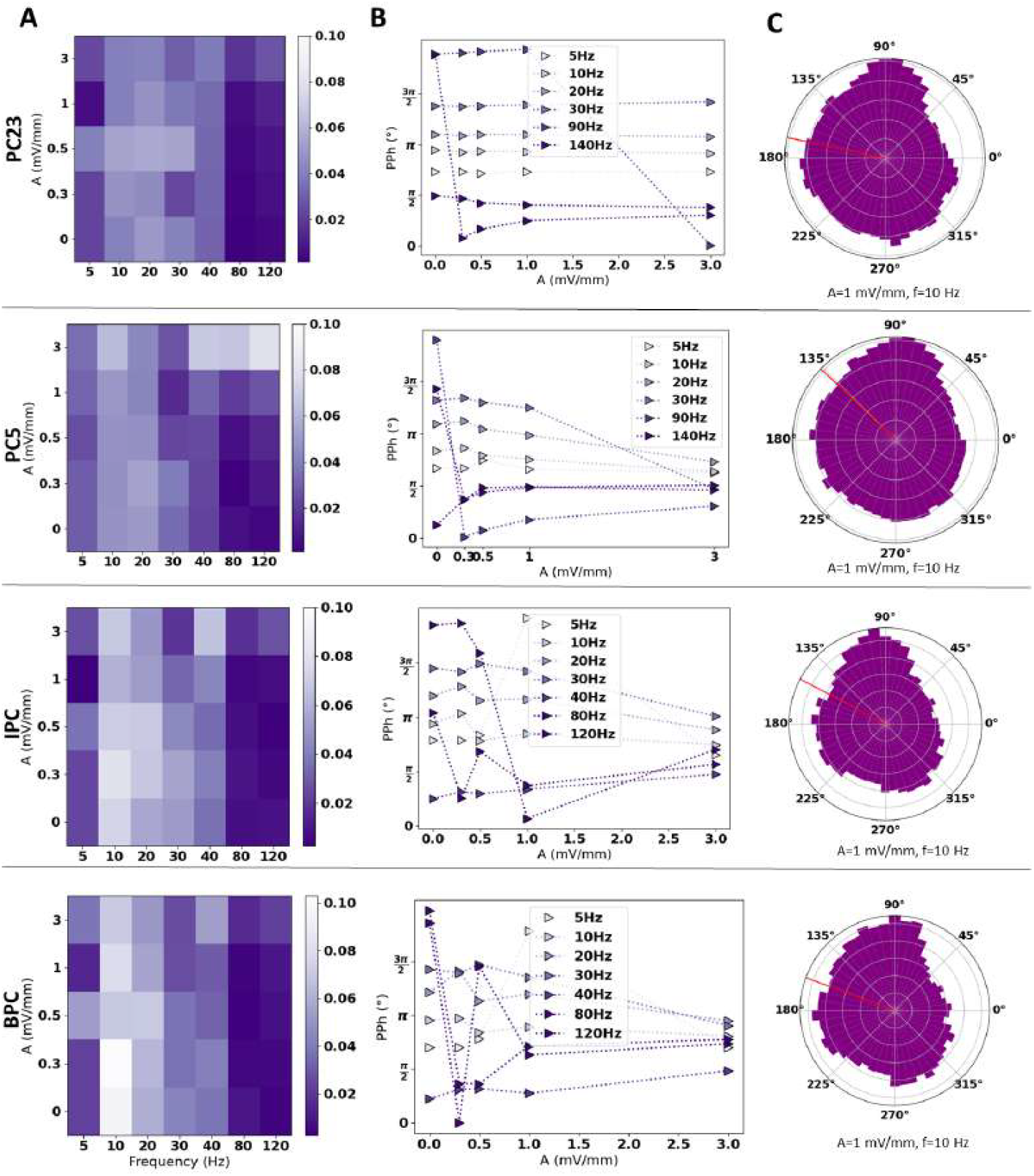
Impact of tACS on the network activity of Principal Cells (PC) of layer 23 (PC23), layer 5 (PC5), Inverted PCs of layer 6 (IPC) and Bipolar PCs of layer 6 (BPC) on 8Hz alpha rhythm. (A) Entrainment maps of PCs to tACS with varying intensity and frequency. (B) The Preferred Phase (PPh) variation with tACS parameters. (C) An example of the entrainment of the PC’s action potential for 1mV/mm, 10Hz tACS.

**S7 Fig.**
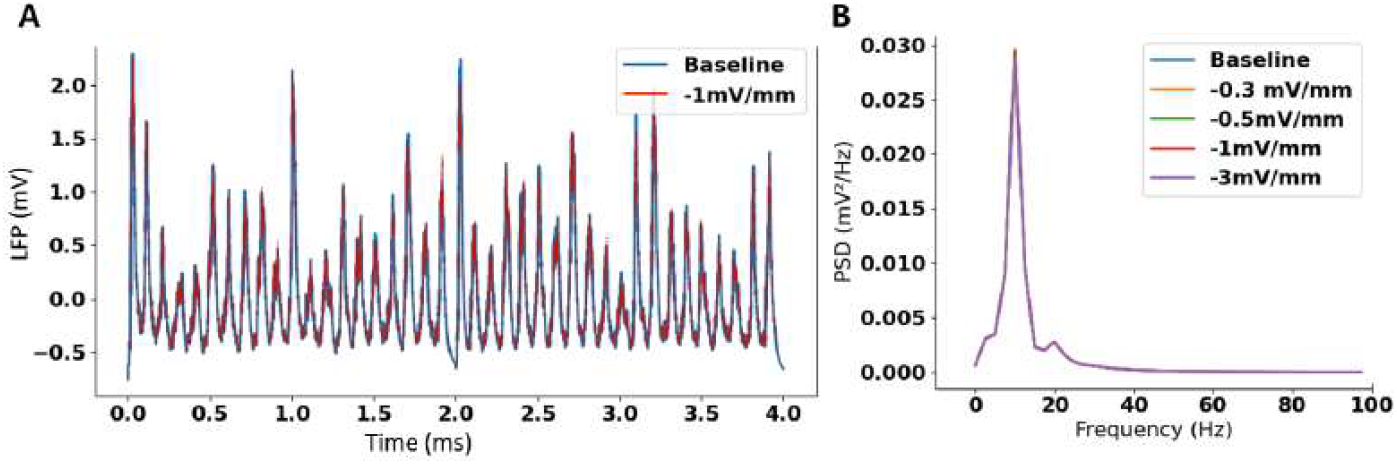
tDCS results for alpha rhythm. (A) An example for the temporal signal at baseline and with 1*mV/mm* cathodal tDCS. (B) Power Spectral Density (PSD) for cathodal tDCS for baseline at 0.3, 0.5, 1, and 3*mV/mm*.

**S1 Table.** The subtypes, e-types, and morphology names of the cells from the library of Blue Brain models used to compute the polarisation lengths used in this study. PC: Principal Cells, TTPC: Tufted Pyramidal cells, UTPC: Untufted Pyramidal cells, IPC: Inverted Pyramidal Cells, BPC: Bipolar Pyramidal Cells, LBC: Large Basket cells, MC: Martinotti Cells, BP:Bipolar Cells, NGF: Neurogliaform Cells, PYR: Pyramidal cells, continuous; d, delayed; b, bursting. AC, accommodating; NAC, non-accommodating; STUT, stuttering, IR, irregular; AD, adapting.

| Layer | Cell type | Subtype | e-type | morphology |
| --- | --- | --- | --- | --- |
| L23 | PC | PYR | cAD | pyr229 (4) |
|  | PV+ | LBC | dNAC | 222 (0) |
|  | SST+ | MC | cACint | 209 (0) |
|  | VIP+ | BP | cACint | 209 (0) |
|  | RLN+ | NGC (DA) | cNAC | 187 (4) |
| L4 | PC | PYR | cAD | pyr230 (1) |
|  | PV+ | LBC | dNAC | 222 (2) |
|  | SST+ | MC | cACint | 209 (0) |
|  | VIP+ | BP | cACint | 209 (0) |
|  | RLN+ | NGC | cNAC | 187 (0) |
| L5 | PC | TTPC2 | cAD | pyr232 (1) |
|  | PV+ | LBC | dSTUT | 214 (0) |
|  | SST+ | MC | cACint | 209 (2) |
|  | VIP+ | BP | cACint | 209 (3) |
| L6 | PC | TPC (L4) | cAD | pyr231 (5) |
|  |  | IPC | cAD | pyr231 (2) |
|  |  | BPC | cAD | pyr231 (4) |
|  | PV+ | LBC | bSTUT | 213 (0) |
|  | SST+ | MC | bIR | 215 (6) |
|  | VIP+ | BP | cACint | 209 (0) |

**S2 Table.** The parameters altered in the NeoCoMM software interface to obtain the simulation types in the manuscript (Figure 2). DC: Distant Cortex, Th: Thalamus, *I*_*inject*_: Injected current density, PC: Principal Cells

|  | Alpha simulations | Gamma simulations | Epileptic signals |
| --- | --- | --- | --- |
| Period (ms) | 100 | 25 | 100 |
| DC Jitter (ms) | 30 | 6 | 10 |
| DC stim number/cell | 13 | 13 | 10 |
| DC $I_{inject}$ ($\mu A/ms$) | 66 | 50 | 60 |
| Th $I_{inject}$ ($\mu A/ms$) | 30 | 30 | 20 |
| Th stim number/cell | 139 | 139 | 20 |
| PC $g_{GABA}$ ($\mu S/ms$) | 0.85 | 0.85 | 25 |
| PC $g_{AMPA}$ ($\mu S/ms$) | 15 | 20 | 10 |

## Acknowledgments

This work has received funding from the European Research Council (ERC) under the European Union’s Horizon 2020 research and innovation programme (grant agreement No 855109).

